# Ethyl Maltol Disrupts Iron Homeostasis in SH-SY5Y Neuroblastoma Cell Line

**DOI:** 10.1101/2023.01.02.522514

**Authors:** Yuchen Sun, Joseph Bressler

## Abstract

Ethyl Maltol (EM) is a commonly used flavoring compound and has been reported to bind iron and facilitate iron transport. Because EM is highly lipophilic, the potential that it disrupts intracellular iron homeostasis was investigated. EM increased the labile iron pool in SH-SY5Y cells and increased iron-responsive protein activity using a reporter assay in the HEK293 cells. EM induced the expression of transferrin receptor 1 mRNA and decreased expression of ferritin light chain protein in SH-SY5Y cells. Expression of the iron-responsive protein amyloid precursor protein attenuated the effects of EM on these iron-responsive genes. EM treatment decreased cell viability and increased DNA damage. EM also induced the phosphorylation of p53 and the expression of the p53 regulated genes, p21 and 14-3-3σ. The expression of APP attenuated the effects of EM on viability, DNA damage, and the p53 response. Overall, we suggest that EM decreases cell viability through a mechanism involving the p53 pathway. The attenuated responses observed in cells expressing APP suggests that the effects of EM are due to disrupting iron homeostasis.

## 1. Introduction

Ethyl maltol (EM, C7H8O3) is widely used as an aromatic additive in food, beverages, pharmaceutical products, and a common additive to vaping fluid used in electronic cigarettes. A study reported that EM is found in 80% of the vaping fluid products survey (Behar, Luo, McWhirter, Pankow, & Talbot, 2018). The FDA classifies EM as “generally recognized as safe (GRAS)” if used at recommended levels, although the guidelines were based on very limited research (Hua et al., 2019). Several studies have shown that EM forms a complex with iron and facilitates iron transport across intestinal membranes (Li et al., 2017). Our group previously reported co-exposure to EM and iron in a cell culture model decreased viability (Durrani, El Din, Sun, Rule, & Bressler, 2021).

Considering EM is highly lipophilic and volatile, EM could easily pass through the blood-brain barrier and enter the brain, which raises the possibility that exposure to EM is neurotoxic by modifying intracellular iron homeostasis. Iron is required in multiple cellular functions including mitochondrial respiration, chromatin modeling, and xenobiotic metabolism (Ke & Qian, 2007). Because the brain is an energy-demanding organ, iron homeostasis is especially important. When iron homeostasis is disrupted, enzymes requiring iron might not be active and reduced iron could cause oxidative damage by generating reactive oxygen species. Indeed, iron and other metals are thought to be involved in the pathologies of Alzheimer’s and Parkinson’s Diseases (Barnham & Bush, 2014).

The homeostasis of iron in cells is tightly regulated. A labile iron pool (LIP) is available for the proteins that require iron for activity (Philpott, Patel, & Protchenko, 2020). Transferrin receptor 1 (TfR1) and divalent metal transporter-1 regulate iron uptake, while ferroportin (FPN) mediates iron export and ferritin light chain protein (FTL) and heavy chain protein (FHP) are involved in iron storage (Ke & Qian, 2007). The mRNAs of the transporter genes have iron response elements (IRE) at the 3’ untranslated regions and the mRNAs of FPN and FLP and FHP have IREs at the 5’ end. When cells are iron-deficient, iron-responsive proteins (IRPs) bind to the IRE resulting in DMT1 and TfR mRNA stability and more iron is brought into the cell. When iron levels are sufficient, FTL and FHP, and FPN protein are synthesized, thereby decreasing the levels of iron availability (Ke & Qian, 2007). APP mRNA also contains an IRE element in the 5’UTR that binds to IRPs (Silva & Faustino, 2015). An increase in the levels of APP protein was reported during iron influx due to increased translation of APP mRNA (Duce et al., 2010). Consequently, APP might have a protective role under circumstances when iron homeostasis is disrupted (Tsatsanis, Dickens, Kwok, Wong, & Duce, 2019).

In this study, we examined the effects of EM on iron homeostasis, which contrast with other studies that assessed the combination of EM and metal. We report that EM is cytotoxic and disrupts iron homeostasis in the SH-SY5Y neuroblastoma cell line. Ectopic expression of APP protects cells from the effects of EM.

## 2. Materials and methods

### 2.1 Reagents

Transient transfection was performed with pEGFP-n1-APP (69924, Addgene) using the jetOPTIMUS transfection reagent (Polyplus, Illkrich, France).

Immunofluorescent imaging was performed with the following antibodies: 1) Anti-phospho-Histone γH2A.x (Ser139) Antibody, mouse monoclonal, clone JBW301 (MilliporeSigma, Burlington, MA) 2) goat anti-mouse secondary antibody (Alexa-Fluor 488). Nuclei were stained with ProLong® Gold Antifade agent (Cell Signaling Technology, Danvers, MA). Western blots were performed with the following antibodies: rabbit monoclonal anti-FTH1 (#3998) (Cell Signaling), rabbit Phospho-p53 (Ser15) Antibody #9284 (Cell Signaling), mouse monoclonal anti-β-Actin (Millipore Sigma), IRDye® 680 goat anti-mouse IgG secondary, and IRDye® 800 goat anti-rabbit IgG secondary (Li-cor, Lincoln, NE). Ethyl maltol stock was purchased from Millipore Sigma and resuspended in DMSO for cell culture use. Intracellular iron measurement was performed using the Calcein AM assay kit (4892-010-K, R&D Systems, Minneapolis, MN).

### 2.2 Cell culture

The human neuroblastoma SH-SY5Y cell line was originally obtained from the ATCC. Cells were grown in Dulbecco Modified Eagle Medium (DMEM) high glucose with 10% fetal bovine serum (FBS). Cells were fed with fresh medium three times a week and passaged in 1:4 ratio with 0.25% trypsin when they reach about 80% confluency. The cells were derived from a peripheral neuroblastoma but have been used to model neurons from the central nervous system (Ross, Spengler, & Biedler, 1983). The cell line is used to model catecholaminergic neurons (Kovalevich & Langford, 2013).

### 2.3 Establishing SH-SY5Y-APP cell line

SH-SY5Y cells were plated in 6-well plates at a density of 400,000 cells per well and transfected the following day with pEGFP-n1-APP (69924, Addgene plasmid) using the jetOPTIMUS transfection reagent (Polyplus, Illkrich, France) following protocol provided by the manufacturer. Because the plasmid also encodes neomycin acetyltransferases, cells overexpressing APP were selected by adding 50 µM of neomycin (G418) for 7 days. Resistant cells were grown and cryopreserved. The levels of APP mRNA were almost 5-fold greater in SH-SY5Y-APP cells compared to SH-SY5Y cells (Fig. 1 supplement).

### 2.4 Intracellular iron measurement

The fluorescent dye Calcein-AM is used to measure the relative levels of labile iron pool (LIP) because it binds to labile iron resulting in quenching (Cabantchik, Glickstein, Milgram, & Breuer, 1996) (Kakhlon & Cabantchik, 2002).

SH-SY5Y cells were plated in 24-well plates at a concentration of 50,000 cells per well, and treated with EM or deferoxamine one day after plating. After a 24-hour treatment, the medium was aspirated and Calcein AM (4892-010-K, R&D Systems, Minneapolis, MN) was added to cells at a final concentration of 4µM. Cells were removed with trypsin after a 15-minute incubation with calcein-AM and transferred into 96-well Costar Black Walled plates at 30,000 cells per well. The level of fluorescence was read at 490 nm excitation and 520 nm emission wavelength on Molecular Devices SpectraMax M5 SoftMax Pro 7.0.3.

### 2.5 Iron-responsive protein (IRP) binding to the iron response element (IRE)

The assay uses a bicistronic reporter gene construct in which IRE is placed 5’ from the Yellow Fluorescence Protein (YFP) gene and the Cyan Fluorescence Protein (CFP) gene is constitutively expressed (Nie & Htun, 2006). When cells are iron deficient, the IRP binds to the IRE located at the 5’-of the mRNA resulting in a decrease in YFP mRNA translation, which is measured to an excitation at 513 nm and an emission at 527 nm. CFP fluorescence assesses transfection efficiency and is measured at an excitation of 439 and emission at 476. The ratio of YFP/CFP fluorescence indirectly measures IRP binding to the IRE. Transient transfections were accomplished with the jetOPTIMUS transfection reagent (Polyplus, Illkrich, France) as described above.

### 2.6 Viability Assay

Cells were plated in 96-well plates at a density of 40,000 cells per well and treated with EM at different concentrations for 24 hours after plating.

(3-(4,5-Dimethylthiazol-2-yl)-2,5-diphenyltetrazolium bromide (MTT) was added to the cells to achieve a 1.2mM final concentration. The cells were returned to the incubator, and after 2 hours, MTT was aspirated and 200 µL of dimethyl sulfoxide was added to each well to solubilize the MTT crystals. MTT-formazan absorbance was read at 540 nm. Background absorbance was measured by adding 10% SDS to wells of untreated cells prior to adding MTT. Percent viability was computed by the following formula, which 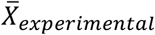 refers to the average absorbance measurement of 6 biological replicates, 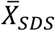 refers to the average absorbance measurement of SDS background and 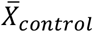 refers to the average absorbance measurement of the medium control.

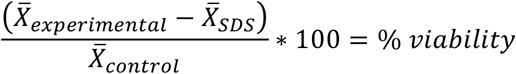

### 2.7 mRNA Measurement

Total RNA was extracted with TRIzol Reagent (Thermo-Fisher, Waltham, MA). Phases were separated with chloroform and the RNA was precipitated with 100% isopropanol. The precipitate containing the total RNA was washed with 75% ethanol to remove impurities. RNA was resuspended in nuclease-free water and the concentration of RNA was quantified with NanoDrop (Thermo-Fisher, Waltham, MA). First-strand cDNA was synthesized from 1µg of total RNA using Oligo(dT)12-18 (500 µg/mL), 10mM dNTP Mix, 5X first-strand Buffer, M-MLV RT reagent purchased from Invitrogen. Primers are described in Table 1 (supplement).

The levels of mRNA were quantified and analyzed using Bio-Rad CFX Manager software to compute the relative quantity (ΔCq) and normalize with housekeeping gene RPLPO to compute the normalized expression (ΔΔCq). The primers sequences for amplification of the target genes can be found in table 1in the supplementary data.

### 2.8 Western blots

Cells were plated at 2,000,000 cells per 10 cm^2^ plate and treated for 24 hours after plating. Cells were collected by scraping into PBS to harvest protein. The cell suspension was pelleted by centrifugation, and protein was extracted using RIPA lysis buffer containing protease and phosphatase inhibitors. Lysates were sonicated and placed on ice for 30 minutes. Soluble protein was isolated by centrifuging lysates at 15,000g for 10 minutes and then harvesting the supernatant. Protein concentration was measured using the Bradford assay.

SDS-PAGE was performed on 4-20% gels subjected to 200 and 400 mA for 40 minutes. Protein was transferred to a nitrocellulose membrane at 200V, 400 mA for 1 hour. The nitrocellulose membrane was incubated with 1X PBS+ 1% Casein Blocker (Bio-Rad) for 1 hour to prevent nonspecific binding. The nitrocellulose membrane was incubated with primary antibodies against ferritin light chain (FTL), phosphorylated p53 (p-p53) both at 1:1000, and β-actin overnight at 4 °C. After washing, the secondary antibody IRDye® 800CW Goat anti-Rabbit IgG (Li-cor) was added for an hour at room temperature. Images were taken with Li-cor Odyssey and band density was analyzed with Image Studio.

### 2.9 Immunostaining for γ-H2AX

γ-H2AX foci indicate sites where DNA double-strand breaks are being repaired. SH-SY5Y cells were plated on poly-L-lysine (PDL)-coated coverslips at 50,000 cells per well in 24-well plates. SH-SY5Y cells were treated with EM for 36 hours. A positive control was accessed with treatment with 3 µM etoposide for 2 hours. SH-SY5Y cells were fixed with 4% formaldehyde for 20 minutes and permeabilized with 0.2% Triton-X-100 for 10 minutes. Cells were blocked with 1% PBS-BSA to prevent nonspecific binding of antibodies. Antibody against γ-H2AX (Ser139) primary antibody (mouse monoclonal, clone JBW301, EMD Millipore; #05-636) (2 µg/mL) was added overnight at 4 °C and fluorescent-labeled Goat anti-Mouse IgG Secondary Antibody was added for 1 hour at room temperature. After washing with PBS, coverslips were mounted onto slides with 4′,6-diamidino-2-phenylindole (DAPI) to stain nuclei and anti-fade reagent. Percentage of cells display foci was computed by dividing the number of cells with 5 foci or more by the total number of cells and multiplying by 100.

### 2.10 Statistical analysis

Graph Pad Prism was used for statistical analysis and making figures. ANOVA and Tukey’s post hos test was used to make multiple comparisons and an unpaired t-test was used determine statistical significance between two groups. The mean of three replicates and the S.E.M. were computed in Prism except for viability assays, where the mean of six replicates was computed.

## 3. Results

### 3.1 EM disrupts iron homeostasis

Iron homeostasis involves regulating LIP so that there is sufficient iron to maintain cellular function. Changes in LIP in either direction would lead to cell death. The LIP was analyzed with the fluorescent dye Calcein-AM. Fluorescence quenching indicates increases in the LIP. Significant decreases were observed in fluorescence in cells at 24-hour after treatment with 0.3mM EM (Fig. 1 A). This concentration of EM is approximately 1/100^th^ the concentration reported in most vaping fluids (Behar et al., 2018). Similarly, a decrease in fluorescence was also observed in cells treated with the iron chelator deferoxamine. Overall, both small molecular weight compounds increase the LIP.

**Figure 1.**
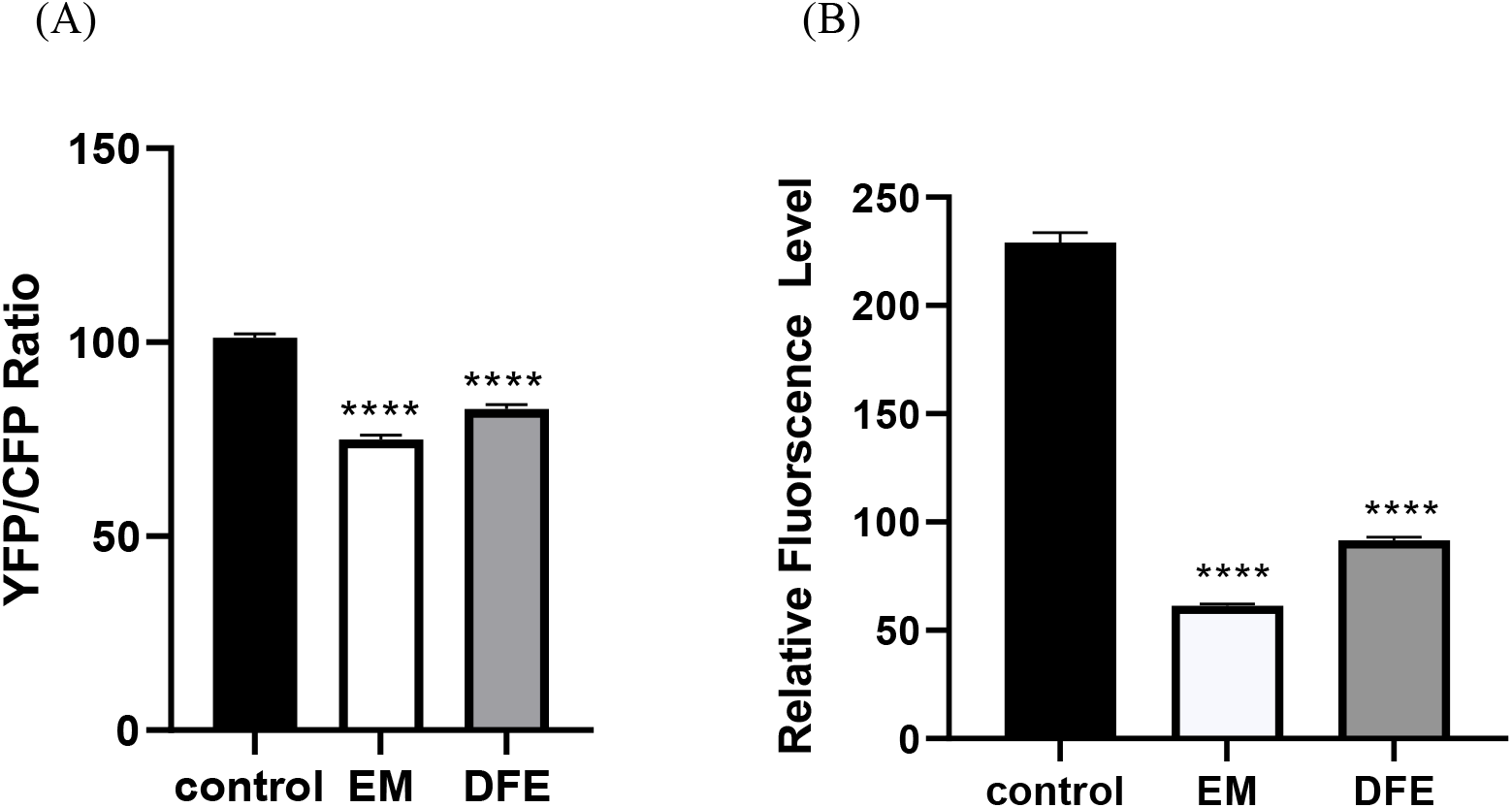
Ethyl maltol disrupts iron homeostasis. (A) Effect of EM on labile iron pool in SH-SY5Y cells. Calcein AM reading was taken at 490 nm excitation and 520nm emission. (B) Effect of EM on iron responsive proteins (IRPs). IRP–IRE interaction in HEK 293 cells after transient transfection of the IRE-containing bicistronic reporter gene. Deferoxamine (DFE), a known iron chelator, was used as the positive control. ****, p<0.0001. (ANOVA, Tukey posthoc test).

To further investigate the effects of EM on iron homeostasis, a reporter assay was performed that indirectly measures the binding of IRP to IRE. The assay was conducted in HEK293 cells to achieve better transfection efficiency. Treatment with EM at 0.3 mM for 24 hours resulted in a significant decrease in YFP/CFP ratio (Fig. 1 B), which indicates greater IRP binding to the IRE (Nie & Htun, 2006).

### 3.2 EM disrupts expression of genes involved in iron homeostasis

An increase in binding of IRP to IRE would result in changes in levels of genes containing an IRE. At 0.3 mM, a 3-fold increase in levels of TfR1 mRNA (Fig. 2 A) was observed in naïve cells. EM did not induce TfR mRNA levels in the SH-SY5Y-APP cell line, however, levels of TfR mRNA were greater in SH-SY5Y-APP cells compared to SH-SY5Ycells.

**Figure 2.**
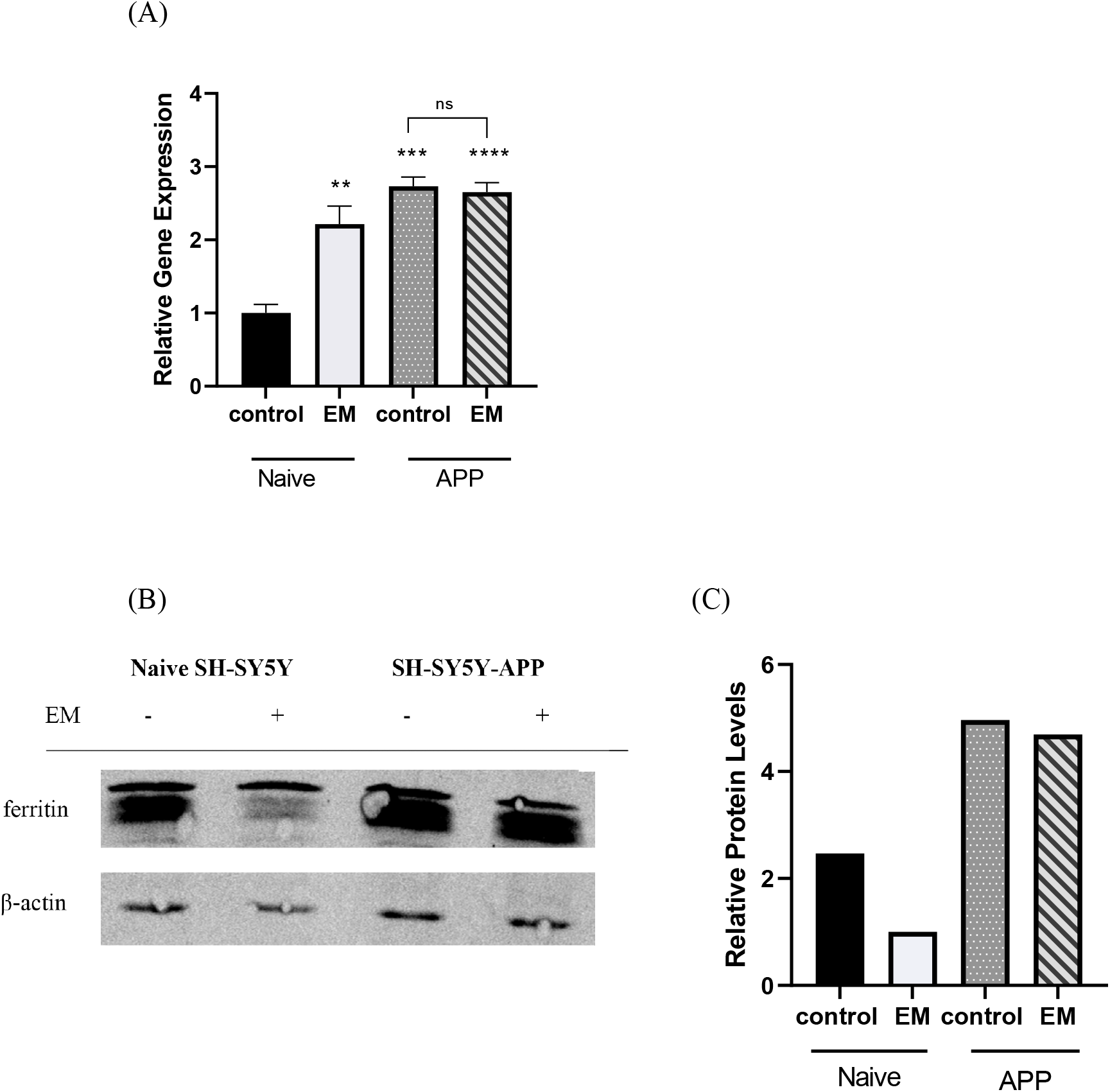
Ethyl maltol modifies expression of genes involved in iron homeostasis. (A) TfR1 levels were measured in SH-SY5Y cells and in APP overexpressing SH-SY5Y cells treated for 24 hr with 0.3 mM EM. TfR1 mRNA levels are normalized to RPLPO using Bio-Rad CFX manager software. Results are expressed as the mean ± S.E.M. (B) Western blot of ferritin in SH-SY5Y cells and in APP overexpressing SH-SY5Y cells after a 24 hr treatment with 0.3 mM EM. (C) Relative protein levels were analyzed with Image Studio and normalized to β-actin (C). **, p<0.05. ***, p<0.01. ****, p<0.0001. ns, not significant. (ANOVA, Tukey posthoc test).

FTL protein was also examined because it has an IRE at the 5’ end. When IRP binds to the IRE, FTL mRNA is not translated resulting in low levels of FTL protein. Interestingly, SH-SY5Y cells treated with EM displayed lower levels of the FTL protein compared to cells not treated (Fig. 2 A). Significant differences were not observed in FTL in APP expressing cells when treated with EM, but the SH-SY5Y-APP cells displayed higher basal levels of FTL (Fig. 2 B and 2 C).

### 3.3 EM treatment decreases cell viability

Chemicals that disrupt iron homeostasis frequently affect cell viability. Approximately a 50% decrease in viability was observed in SH-SY5Y after a 72-hour treatment with 1 mM EM (Fig. 3 A). In contrast, an approximately 25% decrease in viability was observed in SH-SY5Y-APP cells at 1mM EM. These findings provide evidence that EM is cytotoxic to SH-SY5Y cells and the overexpression of APP protects against EM-mediated toxicity.

**Figure 3.**
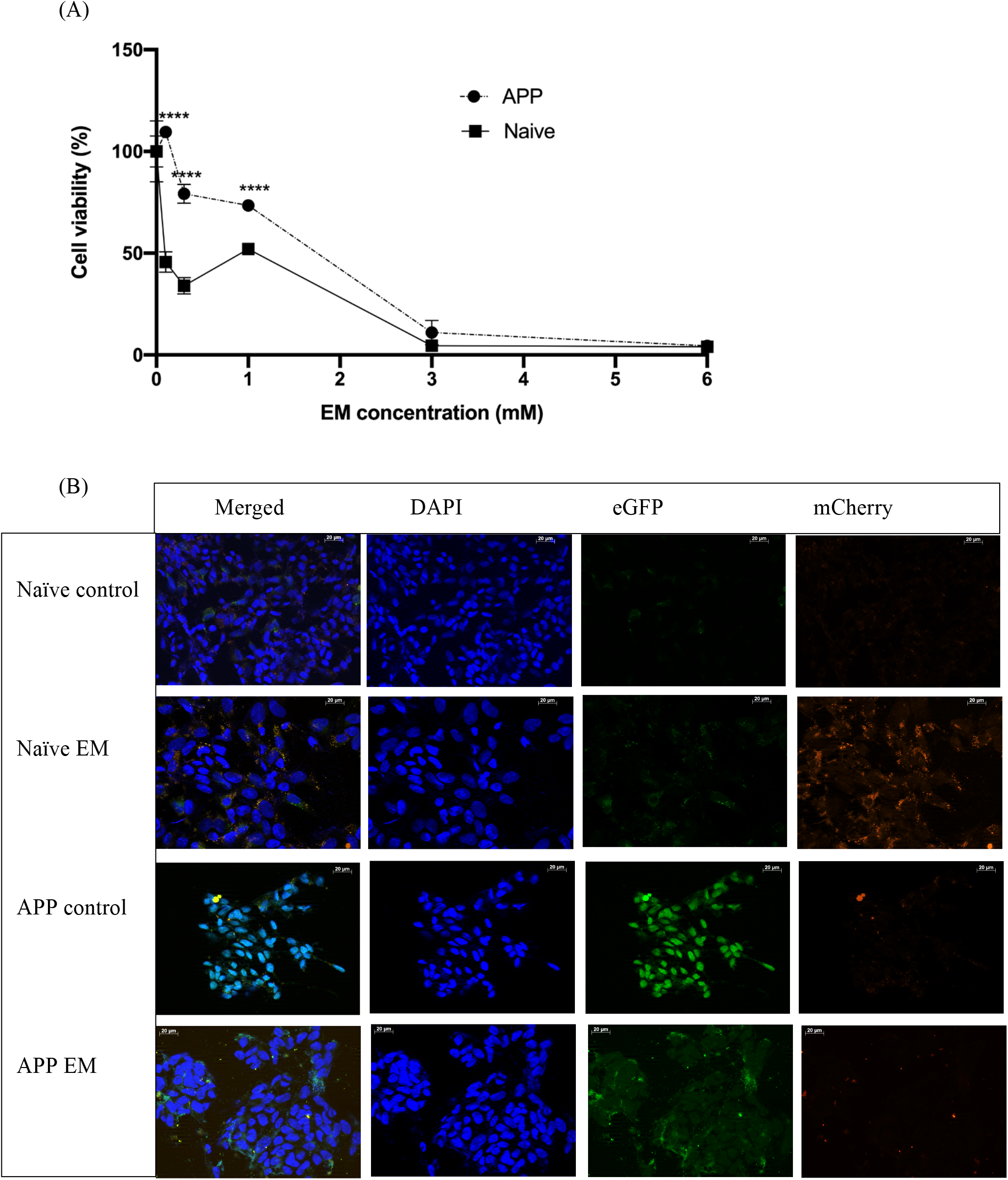
Exposure to EM decreases cell viability and increases DNA damage. (A) MTT assay was performed on SH-SY5Y cell and SH-SY5Y-APP cells that were treated with different concentrations of EM for 72 hours. The dose response curve was generated with Prism 8 in the method of one-way ANOVA and a Tukeys’s HSD was performed to validate the normality of the data. Each value is derived from 6 replicates ± S.E.M. ****, p< 0.0001. (B) γ-H2AX foci immunostaining comparing SH-SY5Y cells and SH-SY5Y-APP cells treated with and without 1mM EM for 72 hours. Cells were fixed and immune-stained with anti-γ-H2AX (mCherry, red) and mounted with medium containing DAPI (blue). APP-positive cells showed green fluorescence in eGFP (green) channel because the plasmd used to express APP was a GFP fusion protein.

In our previous study on EM in lung epithelial cells, the decrease in viability was associated with DNA damage in cells treated with EM. γ-H2AX serves as an early target for DNA damage responses. γ-H2AX foci were not observed in untreated SH-SY5Y naïve cells or SH-SY5 APP cells (Fig. 3 B). At 0.3 mM and 1mM, EM increased γ-H2AX foci formation in both SH-SY5Y cells and SH-SY5Y-APP cells, but the induction was greatly attenuated in the SH-SY5Y-APP cells.

### 3.4 EM induces the p53 pathway

The p53 pathway is activated when cells undergo stress responses such as DNA damage. EM treatment activates the DNA damage response as indicated by the induction of two genes in the p53 pathway, the cell cycle checkpoint genes p21 (Fig. 4 A) and 14-3-3σ (Fig. 4 B). EM induced the expression of 14-3-3σ mRNA in SH-SY5Y cells but not in SH-SY5Y-APP cells (Fig. 4 B). Similarly, EM induced the expression of p21 mRNA in SH-SY5Y cells, which was significant but much smaller in the SH-SY5Y-APP cells. We also examined whether EM treatment increased levels of phosphorylated p53. Phosphorylation increases p53 stability and allows p53-mediated transcription. Higher levels of phosphorylated p53 were observed in SH-SY5Y cells treated with 1mM EM while no significant change was observed in the SH-SY5Y-APP cell after treatment (Fig. 4 C, D)

**Figure 4.**
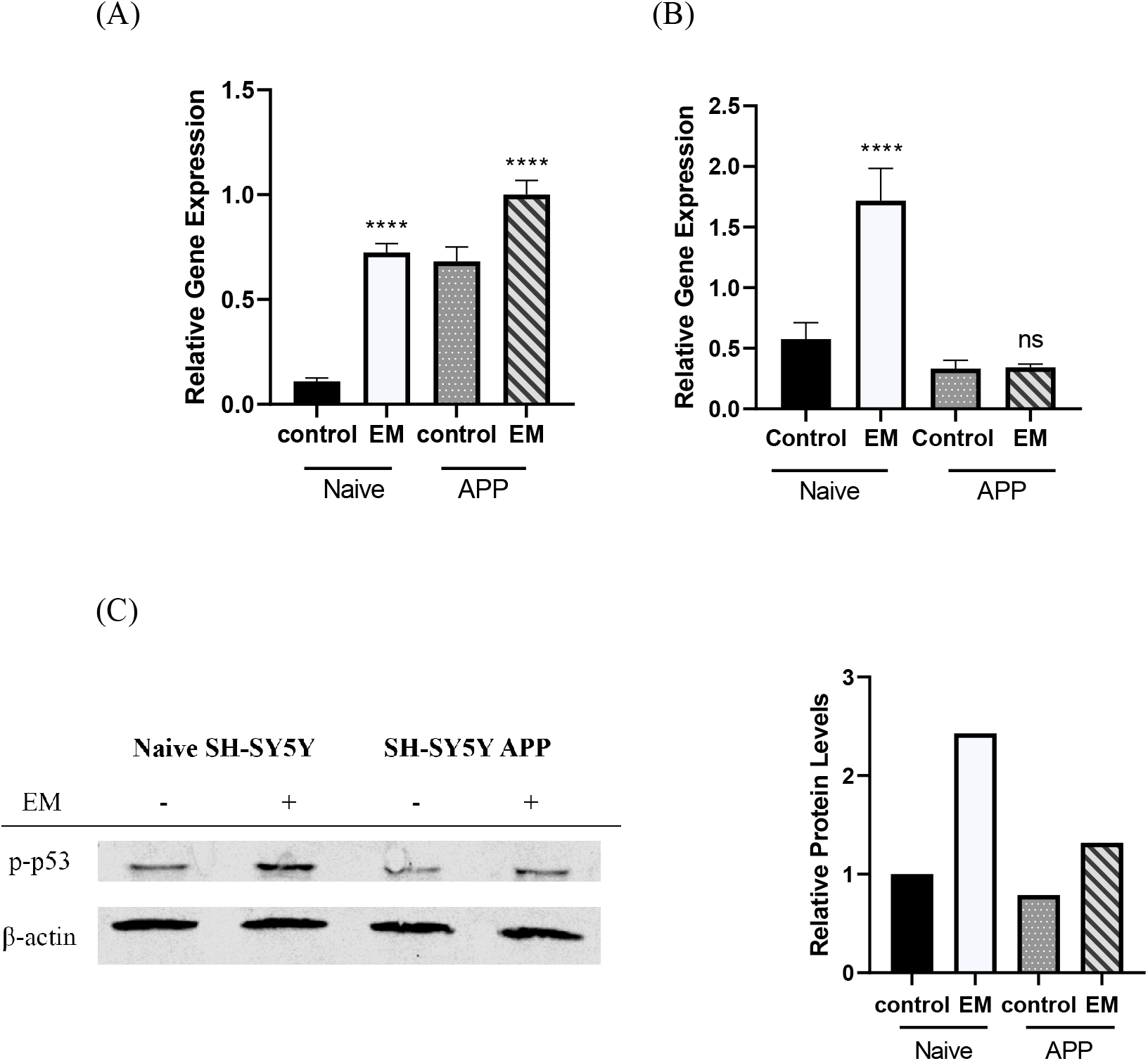
Exposure to EM induces the p53 pathway. (A) Relative p21 gene expression comparing SH-SY5Y cells and APP-SH-SY5Y cells with and without EM at 0.3mM for 72 hours was quantified by measuring levels of mRNA and normalizing to RPLPO by Bio-Rad CFX manager software. (B) Relative 14-3-3σ gene expression was quantified by measuring levels of mRNA and normalizing to RPLPO by Bio-Rad CFX manager software. (C) Western blot was performed with SH-SY5Y cells and SH-SY5Y-APP cells treated with 1 mM EM for 36 hours and blotted with p-p53 and β-actin. Relative protein levels were analyzed with Image Studio and normalized to β-actin. ****, p<0.0001 (ANOVA, Tukey posthoc test).

## 4. Discussion

EM is a lipophilic chemical compound that has the ability to bind and facilitate metal uptake by forming an EM-Fe complex (Li et al., 2017). The complex enters cells resulting in a decrease in viability. Similarly, EM facilitates the effects of copper cell viability in lung cell lines (Durrani et al., 2021).

The question asked here is whether EM disrupts intracellular iron homeostasis and is cytotoxic in the absence of adding extracellular metals. Cells treated with relatively low concentrations of EM displayed increased expression of TfR1 and decreased expression of in FLP, which are indicative of iron deficiency (Silva & Faustino, 2015). Similarly, the increase in IRP activity in the reporter assay is also indicative of iron deficiency. We suggest that EM interacts with iron that is bound to different ligands and creates a pseudo iron deficient state. Interestingly, we also observed higher levels of LIP, which would indicative or iron replete, not iron deficient conditions. It is not clear why the LIP is increasing in cells treated with EM. One possibility is that the higher levels of LIP are due to increased expression of the transporters and lower levels the FTL. We did not measure time dependent changes in LIP, however, and do not know whether increased levels of transporters precede increases in the LIP. Interestingly, in an early study on LIP, the authors observed a reduction in LIP in response to iron chelators followed by a rebound in LIP, which the authors claim was due to mobilization of iron from intracellular iron stores (Konijn et al., 1999). The identity of the iron stores was not investigated. In any event, our studies show that EM modifies iron homeostasis even in the absence of adding free iron to the environment.

Changes in iron homeostasis could explain why EM decreased cell viability. EM might be preventing iron from reaching proteins that need iron for function or excess iron might be killing cells. Both iron depletion and excess has been shown to decrease cell viability (Kasztura et al., 2017). In any event, decreases in viability was likely due to DNA damage. An increase was observed in the percentage of cells that expressed gH2ax, an indicator of DNA damage. Moreover, EM induced activation of the p53 pathway, which initiates the DNA damage response. Increases were observed in phosphorylated p53 as well as two genes that are regulated by p53, namely 14-3 and p21. APP protected the cells against the effects of EM on cell viability and attenuated the DDR. Other studies have also shown APP-mediated protection, for example in models of mitochondrial dysfunction (Cimdins et al., 2019) and mild traumatic injury (Corrigan et al., 2012). Several mechanisms have been suggested to explain APP mediated protection. First, APP might bind iron (Bailey & Kosman, 2019) or APP facilitates ferroportin (FPN1)-dependent iron export at the plasma membrane (Belaidi et al., 2018). In our study, we observed higher levels of FTL and TfR in the SH-SY5Y-APP cell line.

In conclusion, we found that EM decreases cell viability in SH5Y cells and evokes DNA damage. We suggest that the effects of EM are due to disrupting iron homeostasis. EM is highly lipophilic and volatile. Consequently, EM could enter the brain either through the olfactory route or across the blood-brain barrier. Entry through the olfactory route is most concerning because that would bypass biotransformation in the liver and increase risk of disrupting iron homeostasis in the brain.

## Supporting information

Supplemental Table and figuer

## References

Bailey, D. K., & Kosman, D. J. (2019). Is brain iron trafficking part of the physiology of the amyloid precursor protein? J Biol Inorg Chem, 24(8), 1171–1177. doi:10.1007/s00775-019-01684-z

Barnham, K. J., & Bush, A. I. (2014). Biological metals and metal-targeting compounds in major neurodegenerative diseases. Chem Soc Rev, 43(19), 6727–6749. doi:10.1039/c4cs00138a

Behar, R. Z., Luo, W., McWhirter, K. J., Pankow, J. F., & Talbot, P. (2018). Analytical and toxicological evaluation of flavor chemicals in electronic cigarette refill fluids. Sci Rep, 8(1), 8288. doi:10.1038/s41598-018-25575-6

Belaidi, A. A., Gunn, A. P., Wong, B. X., Ayton, S., Appukuttan, A. T., Roberts, B. R., … Bush, A. I. (2018). Marked Age-Related Changes in Brain Iron Homeostasis in Amyloid Protein Precursor Knockout Mice. Neurotherapeutics, 15(4), 1055–1062. doi:10.1007/s13311-018-0656-x

Cabantchik, Z. I., Glickstein, H., Milgram, P., & Breuer, W. (1996). A fluorescence assay for assessing chelation of intracellular iron in a membrane model system and in mammalian cells. Anal Biochem, 233(2), 221–227. doi:10.1006/abio.1996.0032

Cimdins, K., Waugh, H. S., Chrysostomou, V., Lopez Sanchez, M. I. G., Johannsen, V. A., Cook, M. J., … Trounce, I. A. (2019). Amyloid Precursor Protein Mediates Neuronal Protection from Rotenone Toxicity. Mol Neurobiol, 56(8), 5471–5482. doi:10.1007/s12035-018-1460-7

Corrigan, F., Vink, R., Blumbergs, P. C., Masters, C. L., Cappai, R., & van den Heuvel, C. (2012). Characterisation of the effect of knockout of the amyloid precursor protein on outcome following mild traumatic brain injury. Brain Res, 1451, 87–99. doi:10.1016/j.brainres.2012.02.045

Duce, J. A., Tsatsanis, A., Cater, M. A., James, S. A., Robb, E., Wikhe, K., … Bush, A. I. (2010). Iron-export ferroxidase activity of beta-amyloid precursor protein is inhibited by zinc in Alzheimer’s disease. Cell, 142(6), 857–867. doi:10.1016/j.cell.2010.08.014

Durrani, K., El Din, S. A., Sun, Y., Rule, A. M., & Bressler, J. (2021). Ethyl maltol enhances copper mediated cytotoxicity in lung epithelial cells. Toxicol Appl Pharmacol, 410, 115354. doi:10.1016/j.taap.2020.115354

Hua, M., Omaiye, E. E., Luo, W., McWhirter, K. J., Pankow, J. F., & Talbot, P. (2019). Identification of Cytotoxic Flavor Chemicals in Top-Selling Electronic Cigarette Refill Fluids. Sci Rep, 9(1), 2782. doi:10.1038/s41598-019-38978-w

Kakhlon, O., & Cabantchik, Z. I. (2002). The labile iron pool: characterization, measurement, and participation in cellular processes(1). Free Radic Biol Med, 33(8), 1037–1046. doi:10.1016/s0891-5849(02)01006-7

Kasztura, M., Dziegala, M., Kobak, K., Bania, J., Mazur, G., Banasiak, W., … Jankowska, E. A. (2017). Both iron excess and iron depletion impair viability of rat H9C2 cardiomyocytes and L6G8C5 myocytes. Kardiol Pol, 75(3), 267–275. doi:10.5603/KP.a2016.0155

Ke, Y., & Qian, Z. M. (2007). Brain iron metabolism: neurobiology and neurochemistry. Prog Neurobiol, 83(3), 149–173. doi:10.1016/j.pneurobio.2007.07.009

Konijn, A. M., Glickstein, H., Vaisman, B., Meyron-Holtz, E. G., Slotki, I. N., & Cabantchik, Z. I. (1999). The cellular labile iron pool and intracellular ferritin in K562 cells. Blood, 94(6), 2128–2134.

Kovalevich, J., & Langford, D. (2013). Considerations for the use of SH-SY5Y neuroblastoma cells in neurobiology. Methods Mol Biol, 1078, 9–21. doi:10.1007/978-1-62703-640-5_2

Li, Z., Lu, J., Wu, C., Pang, Q., Zhu, Z., Nan, R., … Chen, J. (2017). Toxicity Studies of Ethyl Maltol and Iron Complexes in Mice. Biomed Res Int, 2017, 2640619. doi:10.1155/2017/2640619

Nie, M., & Htun, H. (2006). Different modes and potencies of translational repression by sequence-specific RNA-protein interaction at the 5’-UTR. Nucleic Acids Res, 34(19), 5528–5540. doi:10.1093/nar/gkl584

Philpott, C. C., Patel, S. J., & Protchenko, O. (2020). Management versus miscues in the cytosolic labile iron pool: The varied functions of iron chaperones. Biochim Biophys Acta Mol Cell Res, 1867(11), 118830. doi:10.1016/j.bbamcr.2020.118830

Ross, R. A., Spengler, B. A., & Biedler, J. L. (1983). Coordinate morphological and biochemical interconversion of human neuroblastoma cells. J Natl Cancer Inst, 71(4), 741–747.

Silva, B., & Faustino, P. (2015). An overview of molecular basis of iron metabolism regulation and the associated pathologies. Biochim Biophys Acta, 1852(7), 1347–1359. doi:10.1016/j.bbadis.2015.03.011

Tsatsanis, A., Dickens, S., Kwok, J. C. F., Wong, B. X., & Duce, J. A. (2019). Post Translational Modulation of beta-Amyloid Precursor Protein Trafficking to the Cell Surface Alters Neuronal Iron Homeostasis. Neurochem Res, 44(6), 1367–1374. doi:10.1007/s11064-019-02747-y

