## Supplemental Table and figuer for "Ethyl Maltol Disrupts Iron Homeostasis in SH-SY5Y Neuroblastoma Cell Line"

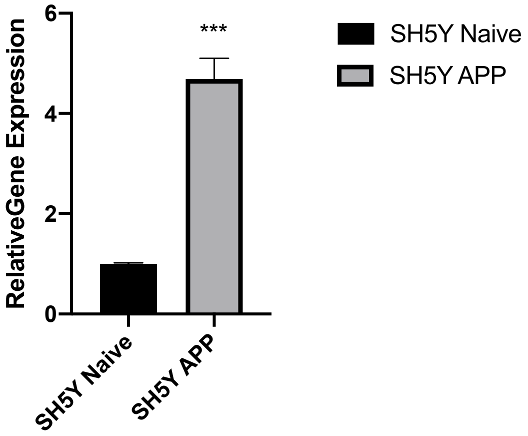


Supplement figure 1. Relative expression of APP mRNA level comparing Naïve SH-SY5Y cells and selected SH-SY5Y APP cells. Relative gene expression was quantified by normalizing to housekeeping gene RPLPO by Bio-Rad CFX manager software. Results are expressed as the mean ± S.E.M. Unpaired T-test was performed to test the significance between two groups. F test was performed to compare variances. ***, p<0.01 *(One-sample t-test)*.

Table 1. Revers and forward primers used for RT-PCR.

| Gene names | Forward 5’ - 3’ | Reverse 3’ - 5’ |
|---|---|---|
| RPLPO | GCAGCATCTACACTGAAG | CACTGGCAACATTGCGGAC |
| APP | GGCCCTGGAGAACTACATCA | AATCACACGGAGGTGTGTCA |
| p21 | TGGACCTGGAGACTCTCAGG | TCCAGGACTGCAGGCTTCCT |
| 14-3-3σ | GGCCATGGACATCAGCAAGAA | CGAAAGTGGTCTTGGCCAGAG |
| TfR1 | ACCATTGTCATATACCCGGTTCA | CAATAGCCCAAGTAGCCAATCAT |
